# Deformable Cell-Like Microlasers for Real-Time Mechanical Quantification in Organoids

**DOI:** 10.1101/2023.06.05.543717

**Authors:** Guocheng Fang, Yu-Cheng Chen

## Abstract

Mechanical stress in multicellular environments plays a critical role in a wide range of tissue function and drug delivery. However, conventional methods are incapable of quantifying internal solid stress *in situ*, which is the hallmark of the 3D multicellular mechanical environment. To overcome the limitations, here we developed hollow-core structured microsphere lasers to realize all-optical direct recording of cellular stress in organoids and spheroids with cellular resolution. The deformations of whispering-gallery-mode laser can therefore be transduced into the change of the laser spectrum to reflect deformation within two-hundred nanometers in deep tissue environment. Our findings demonstrate the capability to quantify internal solid stress in different types of human tumor spheroids in real time. We also explored its potential in mechano-responded drug screening. Dynamic monitoring of contractile stress inside human embryonic stem cell-derived cardiac organoids was also obtained. This method may bring new opportunities to mechanobiology with multicellular resolution and accelerate high-throughput drug screening in human organoids.

## Introduction

Mechanical stress is ubiquitous in various levels of biological systems in contact with surrounding environments^1-3^. Likewise, cells interact with their local microenvironment by sensing and generating forces in response to mechanical cues. This pivotal process becomes more complex when single cells expand into multicellular environments, particularly in three-dimensional (3D) multicellular *in vitro* models. Mechanical-dependent behaviours have been widely revealed in neural, muscle, tumor and stem cells^4, 5^. For instance, solid mechanical force and internal fluid pressure in the tumor microenvironment are critical factors for drug delivery and therapy^6-8^. Heart contractile force maintains arterial blood pressure, matching the metabolic demand^9^. Characterizing mechanical stress in 3D spheroids and organoids is the key to understanding various biological phenomena.

Conventional technologies such as atomic force microscopy, magnetic tweezers, micropillars, micropipette aspiration, microcapsules, and traction force microscopy are powerful tools for measuring biomechanical stress by probing or labelling the external surface of cells, interfaces, or extracellular matrixes^10-12^. However, these methods are incapable of directly characterizing internal solid stress, which is the hallmark of the 3D multicellular mechanical environment. To characterize internal solid stress in the tumor, opening methods are usually involved, which requires the slicing of tumors and is heavily limited to macroscale analysis^13^. Optical imaging based on fluorescence provides a non-contact approach to monitoring biological activities and mechanics inside a multicellular environment. Among all, soft microspheres have emerged as powerful sensors to fill the gap of *in situ* and *in vivo* stress characterization in the 3D cellular environments at supra-cellular scales. Soft microspheres formed by liquid droplets or hydrogels have been embedded into multicellular spheroids or tissues, in which internal solid forces are characterized by comparing the deformation of the microsphere through images (initial state and deformed state)^14-20^.

To date, fluorescent-based soft microspheres are the most used approaches, offering a straightforward tool for quantifying multicellular stress through fluorescence microscopy. However, two critical challenges remain for image-based measurements^21^. (1) Optical imaging suffers from the constraint of light penetration in thick objects (>100 µm) and is often masked by strong background noise from highly scattered emissions in 3D cellular environment. Limited by the diffraction of light, measuring minute deformation (1∼2 hundred nanometres) remains a general challenge for most imaging techniques. (2) Scanning and reconstruction of images are required to obtain accurate changes in surface deformation. Image-based approaches are thus restricted by temporal resolution, which sets a great challenge for monitoring fast dynamic processes of cellular forces in deep tissue environments.

In contrast, laser emissions generated from active optical micro-resonators offer a promising solution with signal amplification and narrow linewidth inside biological systems^22-24^. Instead of analysing images, laser emission provides strong and distinctive lasing spectra which can be used for ultrasensitive detection under free space. Subtle changes in the resonating cavity could therefore be amplified and monitored through lasing spectra. In particular, whispering gallery mode (WGM) microlasers have been widely applied in sensing applications due to their strong light-matter interactions at the spherical cavity interface, including cardiac contractile rhythm, protein binding interactions, temperature, and pH^25-28^. WGM microcavities made from various types of materials have been investigated, such as silica, indium gallium phosphide, liquid crystal, polystyrene, and lipid^29, 30^. Despite the great progress in WGM microlasers, detecting tiny biological forces in cellular environment still remains a challenge owing to the requirement for high refractive indexes (to support lasing) but low modulus of material elasticity to sense tiny forces. To date, most materials used for WGM lasers possess relatively high elastic modulus (*e*.*g*., polystyrene bead ∼3.5 GPa), which cannot be employed to detect tiny multicellular forces (several to tens of kPa). On the other hand, soft materials (such as hydrogels alginate, polyacrylamide, agarose, etc.) usually have a low refractive index which does not support laser generation in biological environments.

Here, we describe a new concept to quantify internal cellular stress by introducing “hollow-core” structured microlasers (Fig.1). The microlasers made by poly(lactic-co-glycolic acid) (PLGA) have ultrathin shells, allowing both great deformability and a high refractive index for the Q-factor. Unlike solid WGM microlasers, the proposed ‘soft’ and cell-like microlasers enable sensitive detection of surrounding stress via tiny deformation. The deformation of microlasers could be shown by the free spectrum range (FSR) of the lasing spectrum. Theoretically, deformation changes could achieve a resolution around 50 nm, and the minimum stress that can be resolved is approximately 0.2 kPa. To illustrate its promising applications, we embedded the microlaser mechanosensors into tumor spheroids and human embryonic stem cell (hESC)-induced cardiac organoids for quantifying the internal solid stress and cardiac contractility, respectively.

**Fig 1.**
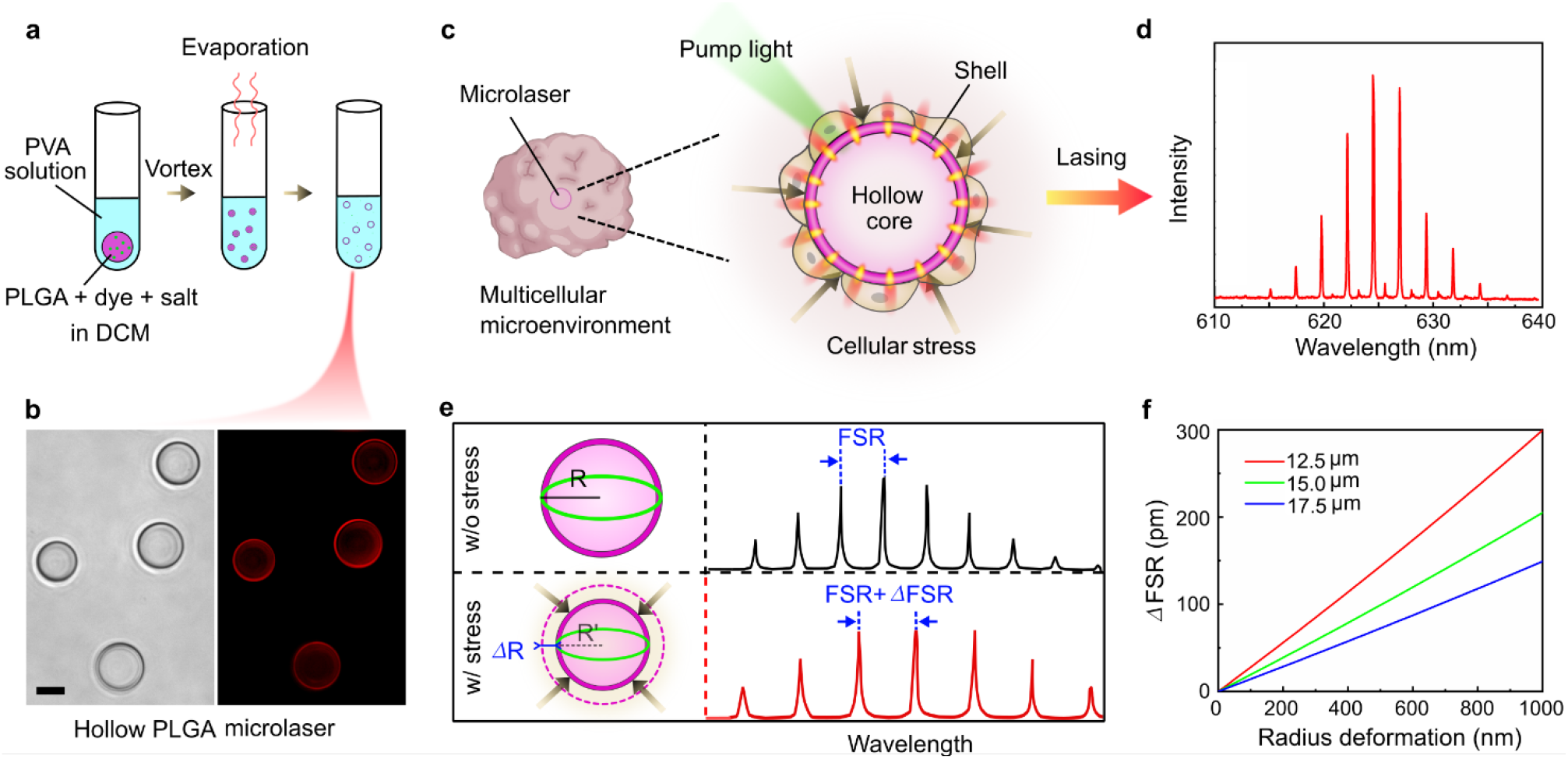
Concept overview of the hollow-core microlasers for cellular stress quantification. (**a**) Fabrication of hollow-core PLGA microlasers. (**b**) Hollow PLGA microlasers under bright-field and fluorescent microscopy. Scale bar: 20 µm. (**c**) Illustration of cellular stress sensing by embedding the microlasers into a multicellular environment. (**d**) A typical laser spectrum generated by a hollow microlaser. (**e**) Stress sensing mechanism by analysing the FSR change: the cellular force (stress) will compress the microbead and reduce the radius, leading to the increase of FSR. (**f**) Relationship of FSR change and radius reduction of three different-size microlasers (under a fixed wavelength of 635nm).

## Results and discussion

### Concept and fabrication of cell-like hollow-core microlaser

Figure 1 illustrates the concept of using hollow-core microsphere laser for mechanical sensing and quantification. To begin with, hollow-core PLGA microlasers were synthesized by employing osmogene-mediated method^31^, as shown in Fig.1a. Details can be found in Methods section and Supplementary Figure 1b. PLGA was selected owing to its good biocompatibility, high refractive index (1.46∼1.49), and low modulus of elasticity^32, 33^. Nile red was doped into the PLGA to serve as the laser gain because of its high lipophilic stability and quantum yield^34^. Figure 1b shows the bright field and fluorescence image of Nile red-doped PLGA microspheres, which have a shell thickness of ∼500 nm.

Figure 1c illustrates the concept of biomechanical sensing by embedding hollow-core microlasers into a multicellular environment (tissues, biological matrix, blood, etc.). Laser emission was obtained under appropriate pump excitation (Fig. 1d). When surrounding cell stress is applied on the microlasers, the microlaser radius reduces accordingly, which thus changes the free spectrum range (FSR) of the laser emission spectrum. Following the formula *FSR=λ*^*2*^*/(2πnR)*, where *λ* is the laser wavelength, *n* is the effective refractive index of the resonator material, *R* is the radius of the microsphere, the radius change of microsphere could be calculated (Fig. 1e). The ultra-narrow laser peak thus enables the precise acquisition of displacement by means of FSR change (ΔFSR). Consequently, multicellular force or stress can be obtained. Fig. 1f gives the relationship between FSR change and radius deformation (displacement) of the microlasers with initial radii 12.5, 15, and 17.5 µm at a fixed wavelength of 635nm. The findings indicated that smaller microlasers could have a higher slope.

### Optical characterization of hollow-core microlasers

Hollow microlasers were imaged under field scanning electron microscopy (SEM) (Fig. 2a). The surface of the microlaser resonators was confirmed to be smooth enough to support WGM resonance (Supplementary Figures 2a-b). An average shell thickness of ∼504 nm could be achieved by adjusting the concentration of NaCl (Fig. 2b, Supplementary Figures 2c-d). We simulated the electric field distribution in a shell when the microlasers (radius: 15 µm, thickness: 500 nm) were immersed in water. The results indicate that this ultrathin shell can support strong optical resonance and offer a theoretical quality-factor of Q=5.30 ×10^4^. By increasing the shell thickness, the Q-factor would slightly increase (Fig. 2c-d); however, the Q-factor of microlasers with different thicknesses remains in a similar order of magnitude. For detecting multicellular stress in organoids, microlasers with 500 nm shells were selected for downstream experiments in this study.

**Fig 2.**
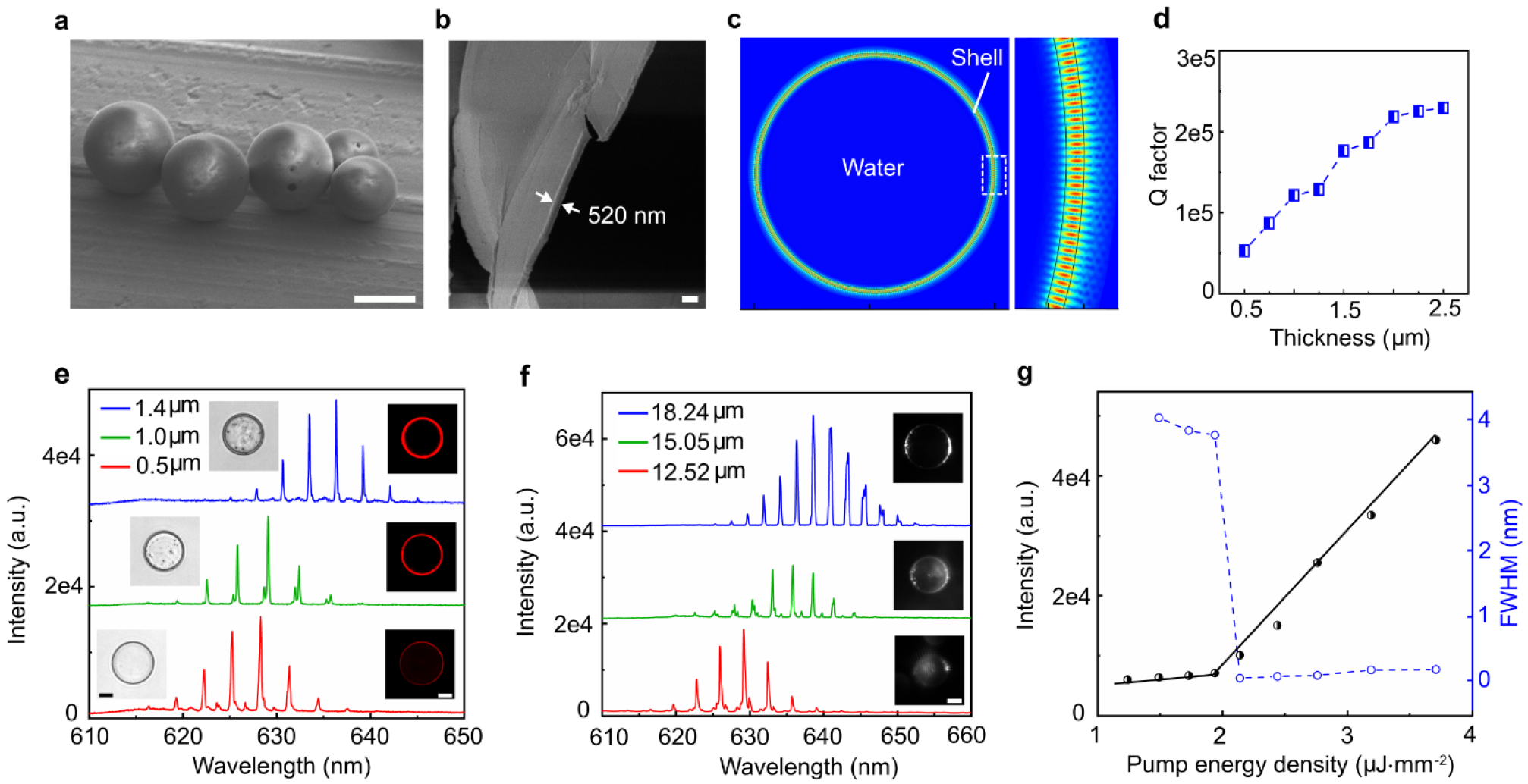
Optical characterization of the Nile red-doped hollow-core microsphere lasers. (**a**) SEM image of the filtered microlasers. Scale bar: 20 µm. (**b**) SEM image of the shell structure. Scale bar: 1 µm. (**c**) Electric filed simulation of a microlaser with the radius of 15 µm. (**d**) Relationship of Q-factor and thickness with the same size (radius 15 µm). (**e**) Laser spectra of the same-size microlasers with thickness of 0.5 µm, 1.0 µm and 1.4 µm, respectively. Inserts are images under bright-field and fluorescent microscopy. Scale bar: 10 µm. (**f**) Emission spectra of microlasers with different radius of 12.52 µm, 15.05 µm and 18.24 µm, respectively. Inserts are images of laser patterns. Scale bar: 10 µm. **g** Spectral integrated intensity of a microlaser with 12.52 µm radius under various pump energy densities. The lasing threshold is ∼2 µJ mm^−2^. The blue curve indicates the FWHM of around 0.033 nm.

Figure 2e shows the lasing spectra of similar-diameter microlasers with thickness of 0.5 µm, 1.0 µm and 1.4 µm excited under the same pump energy density. Insets are the images taken under bright-field and fluorescence microscopy. The emission spectra were measured by an upright microscopy with an excitation wavelength of 552 nm (Supplementary Figure 3). Regardless of shell thickness, the lasing intensity remained in the same order with similar FSR. Microlaser emissions from different diameters were also investigated from 11 to 45 µm, where larger microlasers have smaller FSR (Fig. 2f and Supplementary Figure 4). To verify lasing actions, we measured the output intensities and linewidths by increasing the pump energy densities (Fig. 2g). Spectrally integrated output intensities and linewidths were subsequently measured based on a hollow-core microlaser with a diameter of 12.52 µm. The strong optical feedback of the microlasers greatly reduces the laser threshold to 2 µJ mm^-2^, which is safe for cells and acceptable in biological applications^25, 35-37^. The full width at half-maximum (FWHM) of the laser emission was also measured to be 0.033 nm. Threshold and FWHM analysis for different diameters are provided (Supplementary Figure 5). The emission wavelength of the microlaser can be adjusted by doping other lipophilic organic dyes into PLGA, such as Bodipy-1(Supplementary Figure 6). The microlasers can generate stable laser emission after being stored in water at 4 ºC for up to 2∼3 months. However, PLGA is highly biocompatible and will be degraded eventually^38^.

### Mechanical characterization of hollow-core microlasers

To acquire the mechanical responsivity of the microlaser, we measured the Young’s modulus of the microlasers by atomic force microscopy (AFM) (Fig. 3a). The results indicate that the average Young’s modulus of hollow-core microspheres is approximately 0.62 ± 0.15 MPa (Fig.3b), regardless of the diameter of microsphere. It is worth noting that the Young’s modulus of the PLGA microlasers gradually decreased to 0.60 and 0.46 ± 0.15 MPa after 10 and 30 days, respectively. Yet, laser emissions were still able to be obtained after 30 days (Supplementary Figure 7).

**Fig 3.**
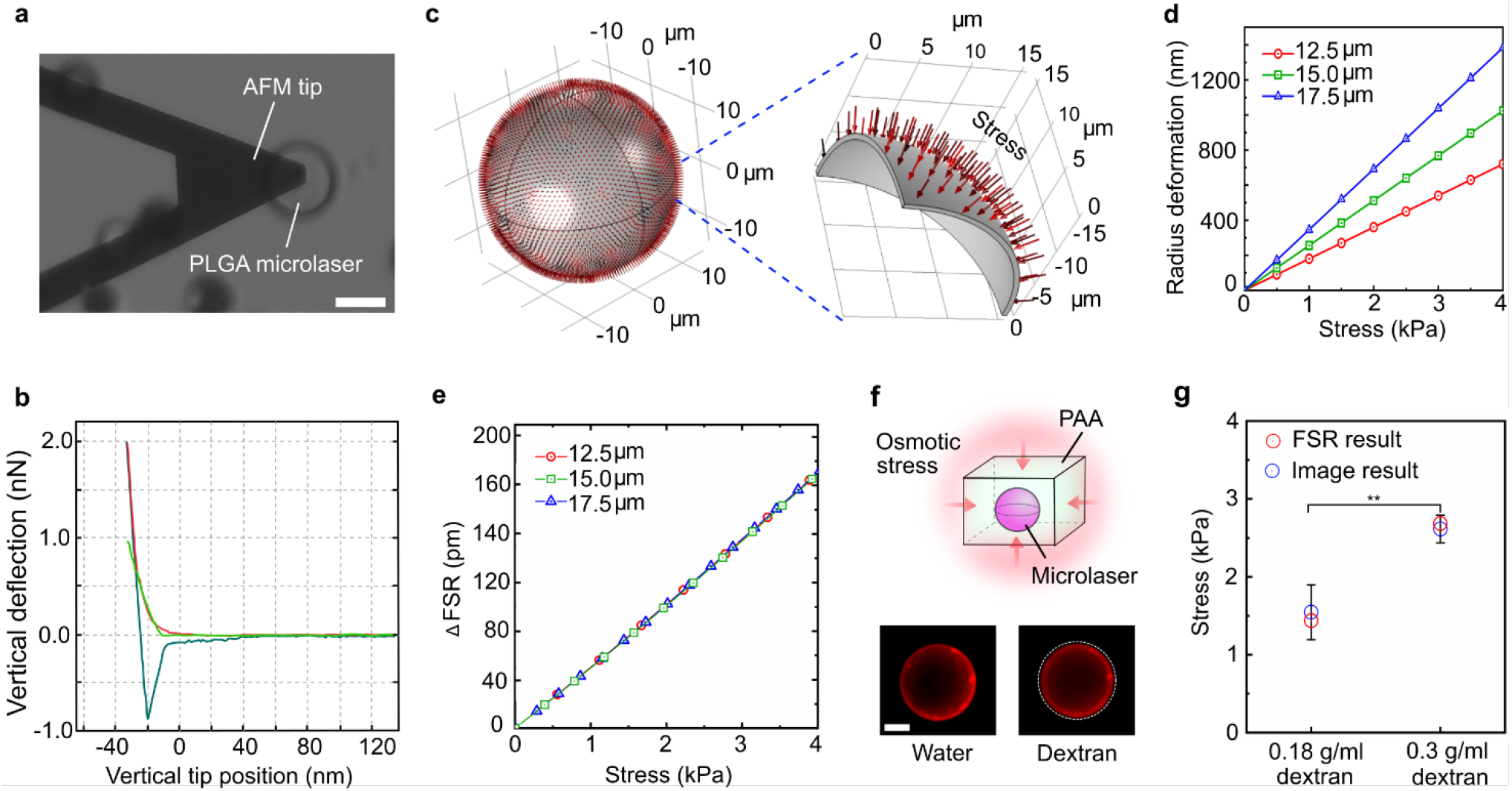
Mechanical characterization of hollow-core microlasers. (**a**) Image of Young’s modulus measurement with AFM. (**b**) Force-distance curve of the microlaser. (**c**) Illustration of the microlaser model suffered from uniform and centripetal stress. (**d**) Relationship of radius deformation and stress. (**e**) Relationship of laser FSR change and stress. Due to the balance of FSR change and radius deformation, the ratios of FSR to stress on different-size microlasers have similar slopes. (**f**) Illustration of microlasers embedded into the PAA hydrogel for the osmotic experiment. The inserts are fluorescent images of a microlaser in water and in 0.3 g/mL dextran. Scale bar: 20 µm. (**g**) Stress calculated by the image-based method and laser FSR-based method (mean ± SD, n=3; *\*\*p*<0.05).

To perform the optical simulation of such structure, here we hypothesized microlaser resonators as a perfect sphere. When they are embedded into the multicellular microenvironment, they will experience uniform and centripetal stress on the surface (Fig. 3c). The mechanical model of hollow elastic microlaser was established and the displacement (in radius) was theoretically calculated (Supplementary Figure 8). We also simulated the deformation and strain using the COMSOL Physics software. The stress distribution of a microlaser (r=15 µm and thickness=500 nm) under 0.6, 1.5 and 2.1 kPa pressure are shown in Supplementary Figure 9. According to the mechanical analysis and simulation, the relationship between radius deformation and applied stress on microlasers with different sizes is shown in Fig. 3d. The larger the sphere, the larger the deformation under the same stress. For any microlaser with a fixed diameter, ΔFSR from lasing spectra should be proportional to the radius deformation (displacement). From Fig.1f and Fig. 3d, the relationship of FSR change and stress were obtained, which indicated that the sensitivities of the three-size microlasers are very close. The average sensitivity is around 51.43 pm/kPa (Fig. 3e, Table 1 and Supplementary Figure 10). Considering the resolution of the spectrum, the minimum detected stress is about 200 Pa. Although this is relatively larger than the stress derived from the single cells (tens of pascal)^39^, it is sufficient enough for the 3D organism stress measurement (several kilopascals to tens of kilopascals)^6, 7^.

To verify the stress derived from spectral-based measurements, we compared the results with conventional image-based measurements. Hollow microlasers were firstly embedded into a bulk polyacrylamide (PAA) hydrogel (Fig. 3f). To induce stress and size reduction of the microlasers, the PAA hydrogel was treated with the dextran solution for the osmotic pressure. Under fluorescence microscopy, we could observe the deformation of the microlasers when the osmotic pressure was applied (Fig. 3f inserts). The results indicated that the deformation of microsphere lasers measured by the image agreed well with that obtained by the laser FSR (Supplementary Figure 11). We further compared the stress calculated by these two methods (Fig. 3g). The measured stress corresponded very well with an error of less than 6.48%. This proves that the accuracy of our proposed method is compatible with that of image-based method. Note that it becomes very challenging for image-based measurements to characterize when the deformation (force-induced displacement) is under submicron scale (Δ <500 nm) owing to the optical diffraction limit.

### Lasing performance in multicellular tumor spheroids

Next, we employed hollow microlasers to detect mechanical stress in the multicellular environment. Multicellular spheroids are excellent 3D models that can fill the gap between 2D cell trials and animal trials. They are formed by the self-assembled cells in suspension, which show higher similarity to native tissues in terms of cellular architecture, cell-microenvironment interactions, metabolism and gene/phenotype^40-42^. The mechanical features of tumor spheroids have a significant effect on tumor growth, metastasis, and anti-tumor drug uptake^43, 44^. Thus, it is highly required to measure the internal stress inside the tumor spheroids.

Here we utilized agarose microwell array to form the spheroids^45^ with a diameter of around 400 µm. To embed hollow-core PLGA microlasers into the multicellular spheroids, we seeded the microlasers into microwells (Fig. 4a), which was followed by cell seeding (Supplementary Figure 12). The quantity of microlaser in each microwell was optimized to be 1∼3. After seeding, the cells gradually aggregated together and grew into compact spheroids due to the cadherin interactions. To improve the cell attachment to the microlasers, the surface of microlasers was coated with fibronectin before coculture with the cells (See Methods). Here we utilized the A549 lung tumor spheroids as the proof-of-concept model (Fig. 4b). The average diameter of the spheroids can be around 260 µm with a seeding concentration of 1.5×10^6^. We stained the nuclei and F-actin of the spheroids and observed them under confocal fluorescence microscopy, confirming the good embedding of the microlasers inside the spheroids (Fig. 4c, Supplementary Figure 13). The cellular structures can still be observed under the microlaser, which may be caused by the optical-lens effect of the transparent hollow-core microspheres^46, 47^. After 3-day coculture, the spheroids remained high viability by Calcine-AM staining, indicating the PLGA microlasers have no toxicity to cells (Supplementary Figure 14).

**Fig 4.**
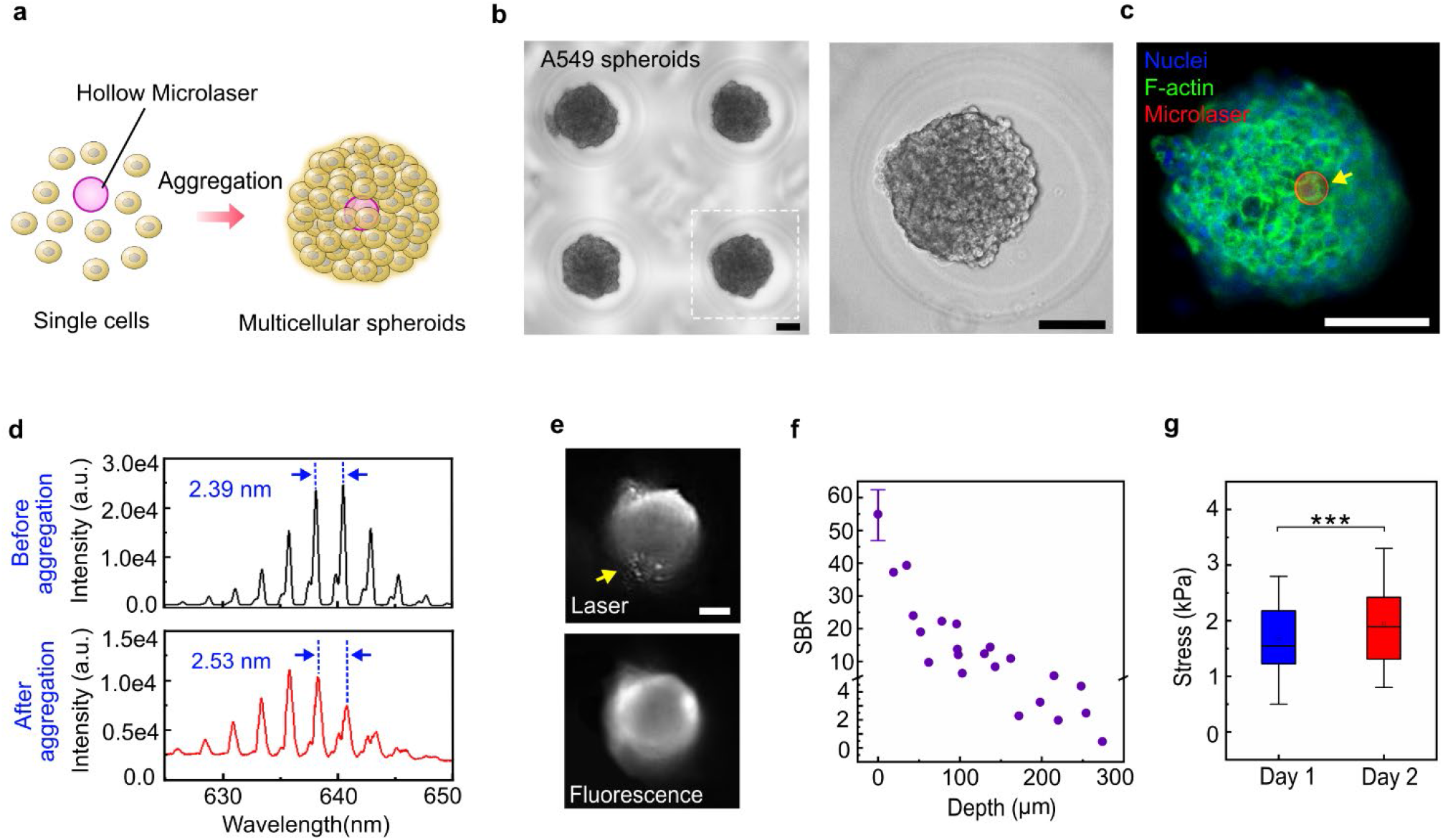
Hollow-core microlasers embedded into multicellular spheroids and their behaviours. (**a**) Illustration of co-culturing the microlaser with cells. (**b**) A549 lung tumor spheroids in a microwell array. Scale bar: 100 µm. (**c**) A microlaser embedded inside an A549 spheroid under confocal microscopy (Blue: nuclei, Green: F-actin and Red: microlaser). Scale bar: 50 µm. (**d**) Emission spectra of a microlaser before and after the cell aggregation. (**e**) Laser image of a microlaser inside the spheroids and its image under fluorescence microscopy. (**f**) Ratio of laser signal to background versus embedding depth. (**g**) Central stress of A549 spheroids measured by the microlaser at day1 and day2 (mean ± SD, n=25, 13; *\*\*\*p*<0.001).

Figure 4d shows the laser emission spectra before and after cell aggregation. The original spectrum presented an FSR of 2.39 nm and increased to 2.53 nm after cell aggregations. Additionally, a slight blue shift was observed after aggregation, which should result from the decreased Q-factor induced by cell attachment to the microlaser surface. Compared to fluorescence emission, we could observe the laser pattern from the laser emission when the microlasers are embedded close to the surface (Fig. 4e, Supplementary Video 1). Due to the significant scattering inside spheroids, the fluorescence emission appears unclear under conventional microscopy. Although cell scattering can decrease the signal intensity and generate background noise, the strong laser emissions still enable us to clearly identify lasing peaks for FSR analysis. The signal-to-background ratio (SBR) acquired from lasing spectral is approximately on the order of few-tens, proving the potential for deep-tissue applications. We then collected the stress at the centre area of the spheroids. It shows that the average stress inside A549 spheroids was around 1.72 kPa on day 1. After one-day growth, the stress increased to 2.02 kPa (Fig. 4g), which agrees well with the reported simulation results^48, 49^.

We are aware that when cells attach to the surface of microlasers, the surrounding refractive index would slightly increase. To test the influence of refractive index on FSR, we measured the FSR of the microlasers in water (RI=1.333), 15 wt% NaCl solution (RI=1.364), and 25%wt NaCl solution (RI=1.378), respectively (Supplementary Figure 15). Our findings show that the refractive index change would not affect the FSR, although the laser peaks showed several-nanometre blueshift. This indicates the refractive index change induced by the cellular microenvironment would not influence the calculation of the stress. We also investigated the influence of temperature ranging from 28ºC to 42ºC. The FSR showed no obvious change, indicating the temperature fluctuation in biological environment has no interference on the stress measurement (Supplementary Figure 16).

### Measurement of solid stress in tumor spheroids

Tumor growth *in vivo* can result in a gradient distribution of oxygen, nutrition, lactate, solid stress, and inertial fluid pressure (IFP)^6, 50, 51^, which can also be observed in tumor spheroids. Here we collected the stress in various depths and gave the stress distribution inside the A549 tumor spheroids (Fig. 5a). The normalized distance (edge to centre) is defined as the ratio of microlaser embedding depth to spheroid radius (Supplementary Figure 17). Given that huge heterogeneity exists among different locations in tumor spheroids, the average stress increased from 1.35 kPa (edge) to 2.01 kPa (centre), with about 49% increment. To test the stress heterogeneity in different tumor spheroids, we selected five kinds of tumor cells (A549, MCF-7, Caco-2, Panc-1, and HepG2) to form the spheroids (lung tumor, breast tumor, colorectal tumor and liver tumor). After one-day culture, solid spheroids with different morphology was formed (Fig. 5b). The average central stress in the spheroids was measured by microlasers. According to Fig. 5c, the central stress in HepG2 liver tumor spheroids could reach up to 2.43 kPa. The lowest stress was observed in Panc-1 pancreatic tumor spheroids with about 1.77 kPa.

**Fig 5.**
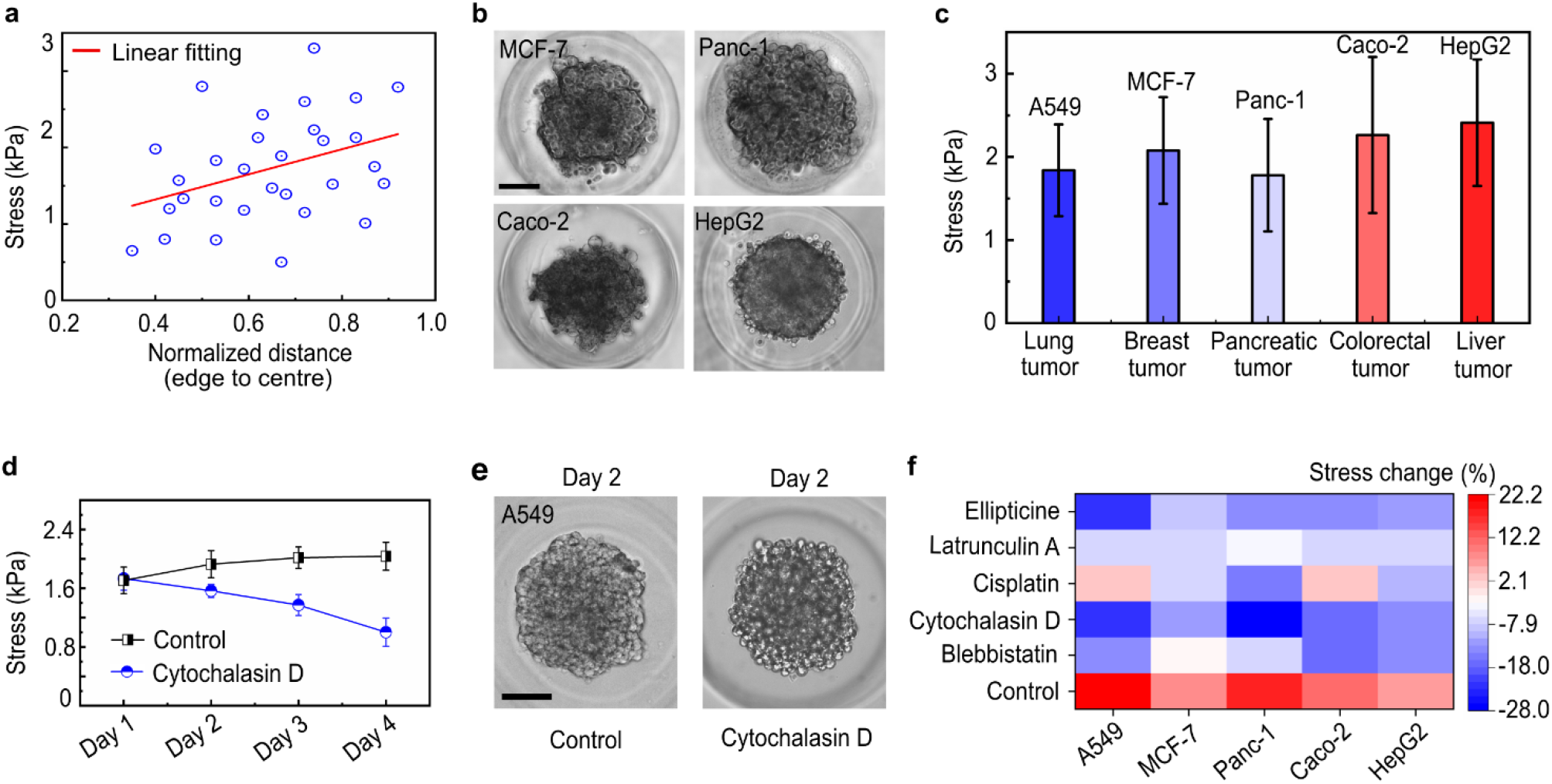
Stress measured by microlasers in tumor spheroids. (**a**) Stress distribution in A549 lung tumor spheroids along the radius. (**b**) Morphology of various tumor spheroids in microwells, including MCF-7 breast tumor, Panc-1 pancreatic tumor, Caco-2 colorectal tumor and HepG2 liver tumor. Scale bar: 100 µm. (**c**) Average central stress in the tumor spheroids after two-day culture. (**d**) Stress change in A549 spheroids after the treatment of Cytochalasin D. (**e**) Morphology of A549 spheroids before and after the treatment of Cytochalasin D at Day 2. Scale bar: 100 µm. (**f**) Tumor stress response to drugs after two-day treatment.

Under the subcellular level, cytoskeletal filaments contribute an important role in stress inside spheroids. To investigate how the cytoskeleton affects the stress, we selected Cytochalasin D to treat the A549 spheroids and tracked the stress within 4 days (Fig. 5d). Cytochalasin D is an inhibitor of actin polymerization. As a control group, the average stress gradually increased from 1.70 kPa (day 1) to 2.03 kPa (day 4). In contrast, the average stress decreased from 1.73 kPa (day1) to 1.01 kPa (day 4) after Cytochalasin D was applied, with about 42% decline. We noticed that some cells became round and started to un-attach from the spheroid (Fig. 5e, Supplementary Figure 18). This may result in stress release inside the spheroids. Combined with the microwell array, we further investigate the stress response to various drugs (Fig. 5f). Different stress increase was observed in the control group, with the largest rise of about 22.2% in A549 group. Apart from Cytochalasin D, Ellipticine and Blebbistatin also showed effective stress inhibition and decrease, especially in A549, Panc-1 and Caco-2 tumors. These findings indicate that the microlasers could potentially serve as a powerful platform for mechano-related drug screening.

### Measurement of contractile stress in hESC-derived cardiac organoids

Finally, here we demonstrate the capability to perform stress detection and quantification in human organoids. hESC-derived cardiac organoids hold extraordinary promise within cardiovascular research, from disease modelling, transplantation, to drug screening^52-54^. Alterations in cardiac contractile stress observed *in vitro* are of physiological relevance, to evaluate contractile function in cardiac organoids as means to characterise disease phenotypes in response to stressors. In addition, the measurement of cardiac contractile force may also be used to determine the maturation status of cardiac organoids^55^. Various methods have been used to quantify the contractile stress in heart tissues and cardiomyocyte spheroids, such as the micropillar method, piezo-electric sensors, magnetic microbead tweezer and image-based fluorescent microspheres^56, 57^. However, it is challenging to achieve conformal contact in organoids with these rigid sensors. Here, we utilized the proposed soft microlasers to measure the contractile stress in the hESC-derived cardiac organoids (Fig. 6a). Cardiac organoids were cultured in two methods, including 96-well plate and agarose microwell array (see Methods). For those grown in a 96-well plate, the diameter of organoids could reach up to 2 mm (Fig. 6b). For those grown in an agarose microwell array, the organoids maintained a diameter of around 300 µm (Fig. 6c). Figure 6d clearly shows that microlasers were successfully embedded into the organoids.

**Fig 6.**
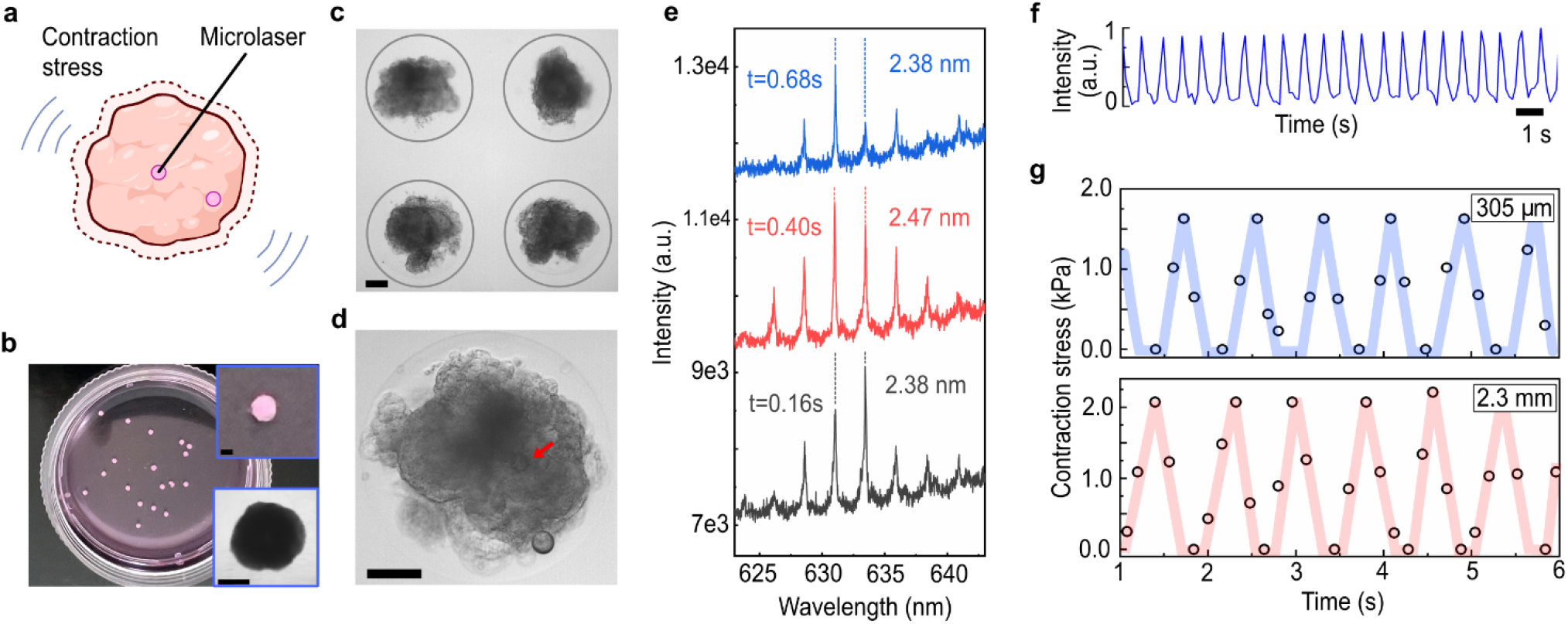
Contractile stress measurement in hESC-induced human cardiac organoids. (**a**) Schematic of the contractile stress measurement in cardiac organoids. (**b**) Image of large cardiac organoids achieved by ultra-low adhesive 96-well plate. Inserts are the enlarged view of the cardiac organoids. Scale bar: 1 mm. (**c**) Small cardiac organoids achieved by agarose microwell array. (**d**) Enlarged view of the small cardiac organoids. Scale bar: 100 µm. Two microlasers could be observed. One is embedded inside the cardiac organoid (marked by a red arrow). Another one is attached to the surface. (**e**) Laser spectra of one microlaser inside a beating organoid. (**f**) Beating frequency (76 bpm) of an organoid with a size of 305 µm. (**g**) Quantified contractile stress of the cardiac organoids with the size of 305 µm and 2.3 mm, respectively.

When the microlasers are located near the surface, the compression can be directly observed in the bright field when cardiac organoids beat (Supplementary Video 2). When embedded in deep tissue locations, the image resolution and contrast becomes a challenging issue (Supplementary Video 3). Here we first show that microlasers could measure recovery actions, returning to their original state after the contraction (Fig. 6e). The lasing spectra of one microlaser were recorded at 0.16, 0.40 and 0.68s. The FSR increased by about 90 pm when the microlaser was compressed. This is consistent with the change in laser spectra. It is worth mentioning that the fast acquisition rate of spectrometer system needs to fulfill the requirement of heart beating frequency (60∼160 bmp).

To demonstrate long-term tracking in human cardiac organoids, we eventually tracked the lasing spectra and analysed the FSR change over several rounds of beating behaviour. To confirm the beating rate of the cardiac organoids, we used fluorescence microscopy as a reference (Fig. 6f). The beating rate of the cultured cardiac organoids ranged from 50 bpm to 120 bpm. According to our measurement, the cardiac organoids with smaller sizes have a contractile stress of about 1.63 kPa. For larger organoids, the contractile stress was approximately 2.07 kPa (Fig. 6g). This value is similar to that of the human stem cell-derived cardiac tissues, but still lower than that of the human adult cardiac tissues^58, 59^.

## Discussion

Whispering-gallery mode laser based on microspheres has been widely applied in biological sensing, not only for its high sensitivity, but most importantly for its compact size (few microns to tens of microns). Compared to fiber-based or Fabry-Perot-based optical resonators, the small size of microsphere microlasers offers the unique opportunity to measure tiny forces inside spheroids and organoids. This is especially important for high-throughput detection and analysis, as the microspheres can be prepared easily and uptake by cells or tumors directly on a massive scale. Such microlasers are also good candidates for deep-tissue localization/sensing as they have distinctive spectral features even at 300 µm depth^27^, which is enough for most applications in spheroids and organoids, compared to imaging-based methods. Combined with microfluidics, this method holds great potential in high-throughput mechanobiology screening on a chip. When integrated with galvo mirror-laser scanning systems, mapping of internal cellular forces could be achieved with multiple resonators inside the organoids.

However, the current technology also presents several limitations and challenges that can be improved. Firstly, to achieve a higher Q-factor and stable lasing, the soft PLGA microlasers must be relatively larger than 20 µm, which is not suitable for single-cell analysis. The size also limits the number of microlasers that can be uptake inside the spheroids and organoids, as they may disturb the stress homeostasis compared to that without microlasers. Searching for high-refractive index materials that can form hollow-core spheres would greatly reduce the resonator size and expand the application. Secondly, the detection resolution of mechanical forces is highly determined by the spectral resolution of the individual spectrometers. The current system is applicable for multicellular forces but insufficient for single-cell forces. To increase the detection capacity, one can measure with a higher grating density or make the microlasers softer by further reducing the shell thickness. Thirdly, the microlaser can only measure the isotropic stress, as the FSR can only reflect the overall diameter change. This is quite different from most image-based fluorescence methods, which can only measure the anisotropic stress^15^. To resolve this issue, the hyperspectral imaging technique may provide the opportunity to analyse laser emissions at different spatial locations with higher spectral precision^30^. This could potentially help the analysis of the anisotropic deformation of the microlasers.

## Methods and materials

### Fabrication of the hollow fluorescent PLGA microlasers

The hollow PLGA microlasers were fabricated by referring to the osmogene-mediated one-step technique^31^. Firstly, 100 mg PLGA (P1941, Sigma-Aldrich) (6.67 % (w/v)) and 5mg Nile red were added to 1.5 mL dichloromethane (DCM). With gentle shaking, the PLGA will be gradually dissolved. Then NaCl crystals were ground by a mortar and a pestle. Then 2 mg NaCl was added to the DCM. Next, the content was stirred for 3.5 h and vortexed for 20 mins to ensure the salt particles distributed in the solution uniformly. Subsequently, the solution was poured into 30 mL 2 wt% polyvinyl alcohol (PVA) aqueous solution in a 50 mL tube, which was followed by the vortex for emulsion. Finally, the solution was poured into a beaker and stirred at 500 rpm and at room temperature overnight to ensure the complete evaporation of DCM. The vortex aqueous-oil emulsion method produced a number of monodispersed microlasers ranging from 5 to 100 µm. After the filter, we achieved microspheres ranging from 20-40 µm. The microlasers smaller than 20 µm cannot generate stable lasers due to their low Q-factors. The achieved microspheres can be washed by reverse osmosis (RO) water and stored in RO water at 4 ºC. The thickness of the microspheres is highly dependent on the concentration of NaCl. We noticed that 2 wt% NaCl can ensure around 500 nm thickness of most microspheres. With the decreasing concentration of NaCl, the thickness of the microspheres will increase. The 1 µm and 1.5 µm microspheres require around 1% and 0.7% concentration.

### SEM and AFM characterization of microspheres

To characterize the morphology and thickness, the microspheres were observed under the SEM (Sirion, FEI Netherlands). Firstly, the microlasers were suspended in RO water and left in an ultrasonic bath for 10 mins to break them. Then the microlasers were loaded on a foil layer and heated for 5 mins at 60 ºC. Finally, they were checked under the SEM.

The Young’s modulus of PLGA was characterized by JPK NanoWizard® 3 NanoScience AFM. The MLCT cantilever (Silicon Nitride Tip A, Bruker’s Microlever AFM probes, k = 0.05 N/m) with a tip radius of 20 nm was selected as the indentation probe. The cantilever sensitivity was calibrated on a hard glass slide and spring constant calibration was performed in a liquid environment using the thermal noise method. The approaching force curves of PLGA spheres were collected in contact mode and further processed by the JPKSPM Data Processing software. After removing the offset and tilt from the baseline, finding the contact point, and subtracting the cantilever bending, the Hertz model was applied to calculate Young’s modulus. The tip was approximated as a paraboloid, and the data range of the fitting was within tens of micrometres.

### Laser spectroscopic characterization

The experimental setup is illustrated in Supplementary Figure 3. All the components were integrated into a Nikon Ni-E upright confocal microscopy. A 10X objective was used for the excitation and signal collection. The pulsed laser (EKSPLA NT230, 50 Hz, 5 ns pulse width) with a parameter oscillator (OPO) was used as an optical pump. The excitation wavelength was set at 552 nm for Nile red, and 448 nm for Bodipy-1. Laser beam size was around 32 µm. The emission laser from microspheres was split by a beam splitter and sent to a spectrometer (Andor Kymera 328i+Newton 970) and a camera (Andor Zyla 5.5) for spectrum and image capture. During the cell laser experiments, the spheroids were kept in an integrated incubator with 37 ºC and proper humidity. The laser threshold was characterized by varying the pump power with a neutral density filter. The SBR is defined as SBR=(*IWGM*-*IBG*)/*IBG*, where *IWGM* is the laser peak intensity and *IBG* is the average background intensity near the laser peak.

### Osmotic experiment

Acrylamide and bisacrylamide (A9099, 146072, Sigma-Aldrich) were mixed and dissolved in water with 5 wt % and 0.11 wt %, respectively. Then the microlasers were mixed with the solution. To polymerize the 1mL of the solution, 10 µL ammonium persulfate (APS, A3678, Sigma-Aldrich 10 % w/v in water) and 0.75 µL N,N,N’,N’-tetramethylethylenediamine (TEMED,T7024 Sigma-Aldrich) was added. Then the solution was vortexed for several seconds and then injected into a mould. After several minutes, the polymerized hydrogel was put into water. To generate the osmotic pressure, 150 kDa dextran was used (D4876, Sigma-Aldrich), which has larger molecules than the pores in PAA hydrogel. Following the osmotic formula, the dextran was dissolved in water with concentrations of 0.18 g/mL (∼ 3kPa) and 0.3 g/mL (∼5 kPa), respectively.

### Cell culture and tumor spheroid formation

Tumor cells (A549, Panc-1, MCF-7, Caco-2 and HepG2) from ATCC were cultured in DMEM medium supplemented with 10% fetal bovine serum (FBS) and 100 U mL^−1^ penicillin/streptomycin. Cells were incubated in a standard incubator with 5% CO2 at 37 °C. When the cells reached 70-80% confluence, a cell passage was performed. The cells could be used after 3-4 passages. We referred to the agarose microwell method to generate spheroids^45^. The agarose chips were fabricated by the 3D-printed moulds (FormLabs Inc. USA). Before loading the cells, the chips were kept in PBS for at least 1 day. Then the cells were treated with Trypsin-EDTA solution for 3 min followed by centrifugation for 5 min at 500 × g. After removing the supernatant, the cells were re-suspended in 1 mL fresh medium and mixed well using a micropipette. The cell concentration was 1 × 10^6^-1 × 10^7^ mL^−1^. Lower concentration will result in smaller spheroids. Finally, the cells were seeded on the microwell chip and remained for 5 mins to ensure sedimentation. They were cultured in the incubator and would form the spheroids after around one-day culture. The medium was changed every 2 days.

### Human embryonic stem cell culture and generation of Hesc-derived cardiac organoids

The human H7 hES cell line were cultured in feeder-free conditions on culture plates coated with Matrigel® matrix (Corning, U.S.A.); maintained in StemMACS™ Ips-Brew XF medium (Miltenyi Biotec, Germany). Culture medium was changed daily. After 80–90% confluency was reached, Hes were passaged using Collagenase Type IV (Gibco, U.S.A.). At 90-95% confluency, differentiation was started using RPMI-1640 Medium (1x) (Hyclone) with B-27 Minus Insulin 50x supplement (Gibco, U.S.A.) and 1% Pen/Strep (Gibco, U.S.A) containing 12µM CHIR99021, a GSK inhibitor (Miltenyi Biotec), as described by Lian et al., 2013^60^. After 24 hours in culture, the medium was replaced with RPMI/B27 minus insulin for 2 days. On day 3, the cells were cultured in the same basal media containing an inhibitor of Wnt signalling, such as IWP-2 (Miltenyi Biotec). On day 5, the medium was replaced with the same basal media. On Day 7, the cells were dissociated into single cells using StemPro-Accutase^®^(Gibco, U.S.A.). The cells were resuspended in cardiosphere growing media (CGM) media containing microbeads. Subsequently, 300k Ipsc-derived cardiomyocytes were seeded into each well of a 96-well plate, ultra-low attachment round bottom plate (CoSTAR, 2022-03-14). The plate was spun down at 1000rpm for 5min and the supernatant was removed from each well. With 2x Matrigel^®^ (Corning, U.S.A.) diluted in DMEM/F12 (Gibco, U.S.A.), 15 µL of Matrigel was added into each well, and spun at 1000rpm for 5min. The 96-well plate was incubated at 37 °C under humidified atmosphere of 5% CO2 for an hour. After one hour, 100 µL of CGM media (B27+ supplement, 17.5 ng/mL Thrombin, 3 ng/mL Cardiotrophin-1, 2 ng/mL EGF, 5 ng/mL FGF-2, 20 ng/mL Wnt-3A, 20 ng/mL EDN1) was added into each well, drop-by-drop. Fresh media was changed every two days.

In terms of the seeding in agarose microwells, the hESC-derived cardiomyocytes with a concentration of 1 × 10^6^ were loaded into the microwells. Before that, the fibronectin-modified microlasers were seeded into the microwells. After the sedimentation, the diluted Matrigel was added, which was followed by incubation at 37 °C for half an hour. Then the culture medium was added for long-time culture as mentioned above. After about 2-day culture, the cells automatically assembled together with the microlasers and grew into a compact microsphere. After about 5-day culture, the cardiac organoids started spontaneously beating.

### Immobilization of microlasers with spheroids

To allow cell attachment, the microlasers need to be immersed in fibronectin solution (10 µg mL^-1^) for at least 2 hrs. Then the microlaser concentration was adjusted to around 1×10^4^ mL^-1^ to ensure each well contained 1-2 microspheres on the chip. After loading the microspheres, the cells were seeded on the chip following the above-mentioned method. After one-day culture, the microlasers were embedded inside the spheroids during the spheroid formation.

### Contractile stress measurement in cardiac organoids

The microlasers were seeded into the microwells together with the cells. We arranged that 1∼2 microlasers were seeded in each agarose microwell and 8∼10 microlasers were seeded in each microwell on 96-well plate. When the cardiac organoids started beating, they were measured under the laser system. Here, the acquisition rate of the spectrometer was 25 Hz. During the measurement, the cardiac organoids were remained in a microscopy incubator that can maintain the temperature at 37 ºC. Low temperature can reduce contractile frequency and stress.

### Spheroid staining and imaging

The spheroids collected from the microwell plate were fixed with 4% paraformaldehyde for 1 h. After washing three times with PBS, the spheroids were treated with 0.5% Triton X-100 at room temperature for 1 h. After washing three times with PBS, the Hoechst 33342 (1:1000) and ActinGreen 488 dye (R37110, Thermo Fisher) were applied and incubated for 1h before observation under the confocal microscopy (Nikon Ni-E upright).

### Statistical Analysis

The data were presented by means ± standard deviation (SD). A two-tailed t-test was used to reveal the statistical difference in Figures 3g &4g with Origin 8.0. A p-value less than 0.05 was considered as significant difference (p < 0.05).

## Acknowledgments

This research is supported by A*STAR under its MTC IRG-Grant (No. M21K2c0106). We would also like to thank the support from Nanyang Presidential Postdoctoral Fellowship.

