## Supplementary material for "Deformable Cell-Like Microlasers for Real-Time Mechanical Quantification in Organoids": SI

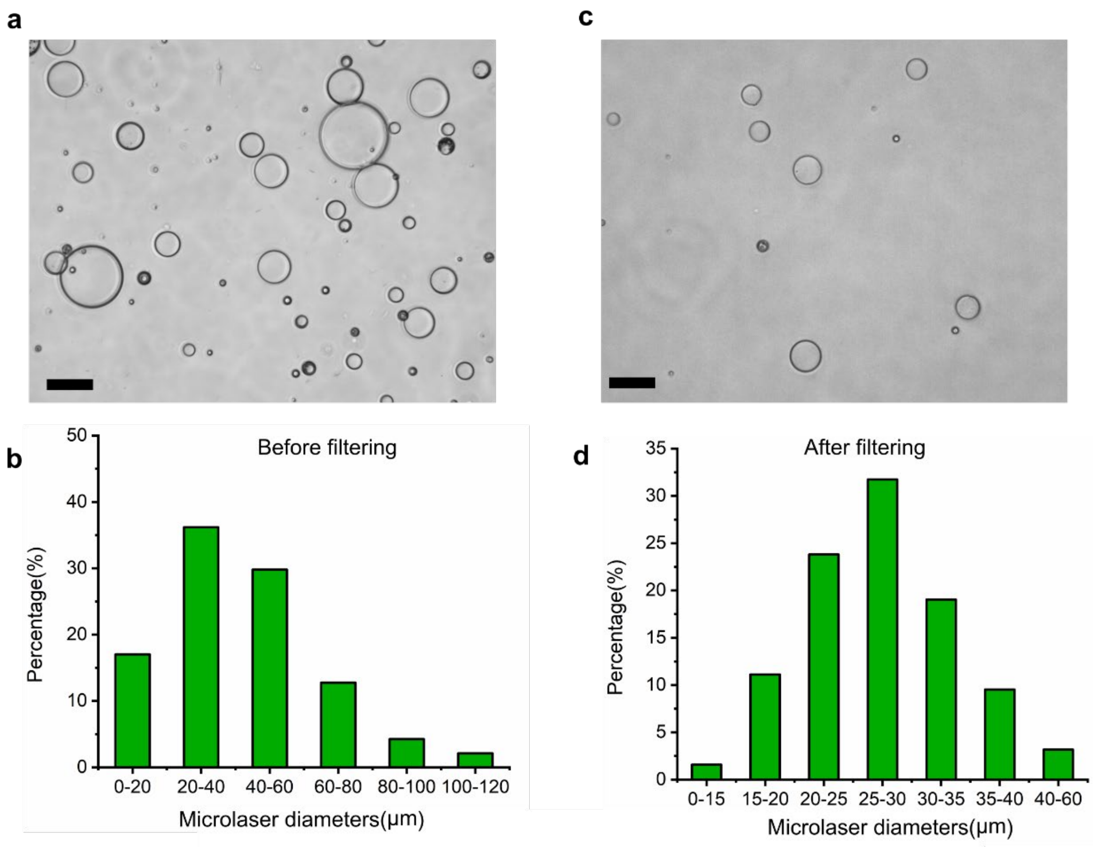


**Supplementary Figure 1**. (a) Image of the hollow PLGA microlaser resonators before filtering. Most of the microlasers are transparent with uniform shell thickness. (b) Size distribution of the hollow-core PLGA microlaser resonators generated by vortex aqueous-oil emulsion method. It can be seen that the candidate microlasers (diameter: 20-40 μm) accounted for around 36.17%. (c) Image of the microlasers after filtering. Scale bar: 50 μm. (d) Size distribution of the microlaser resonators after filtering. We could see that the microlasers ranging from 25 to 35 μm accounted for around 50.79%. This concentration significantly improved the microlaser seeding into the microwell array.

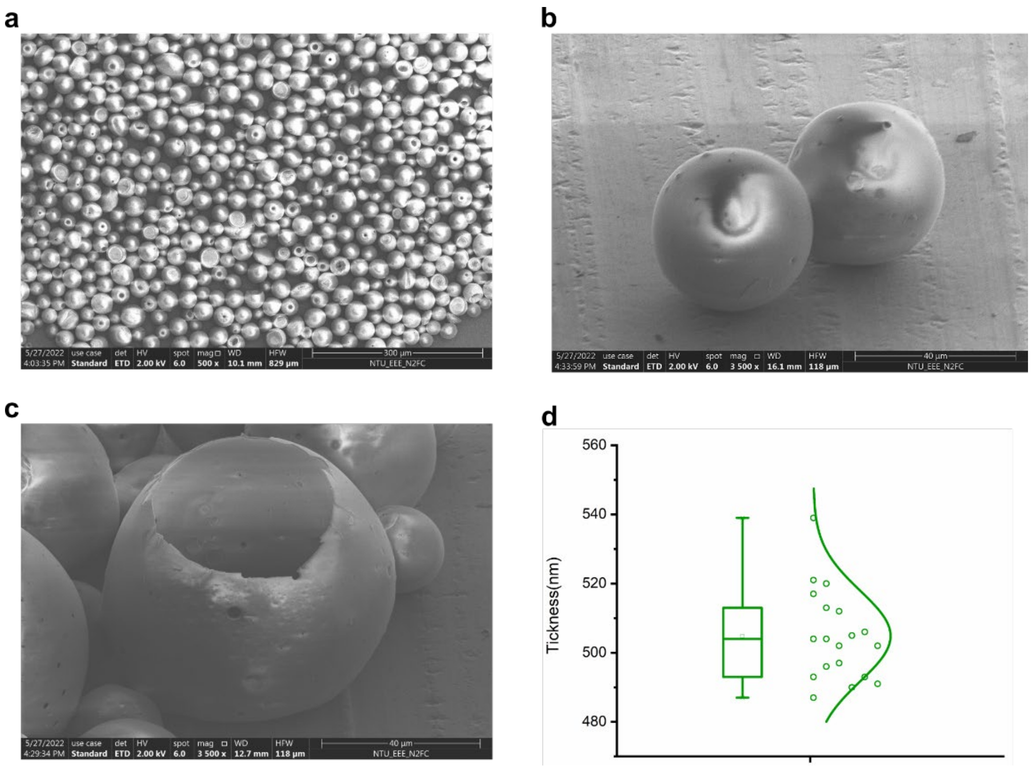


**Supplementary Figure 2**. SEM characterization of the Nile red-doped hollow-core PLGA microlaser resonators. (a) Image of the filtered and dried microlaser. We could generate plenty of microlasers at a time. Some microlaser resonators collapsed due to the ultrathin shell. (b) Enlarged image of the microlaser resonators. We could see the surface of the microlasers is smooth, supporting the light resonating in the cavity. (c) Image of a broken microlaser proving the ultrathin shell. (d) Distribution of the thickness. The average thickness of the microlasers is 504 ± 13 nm.

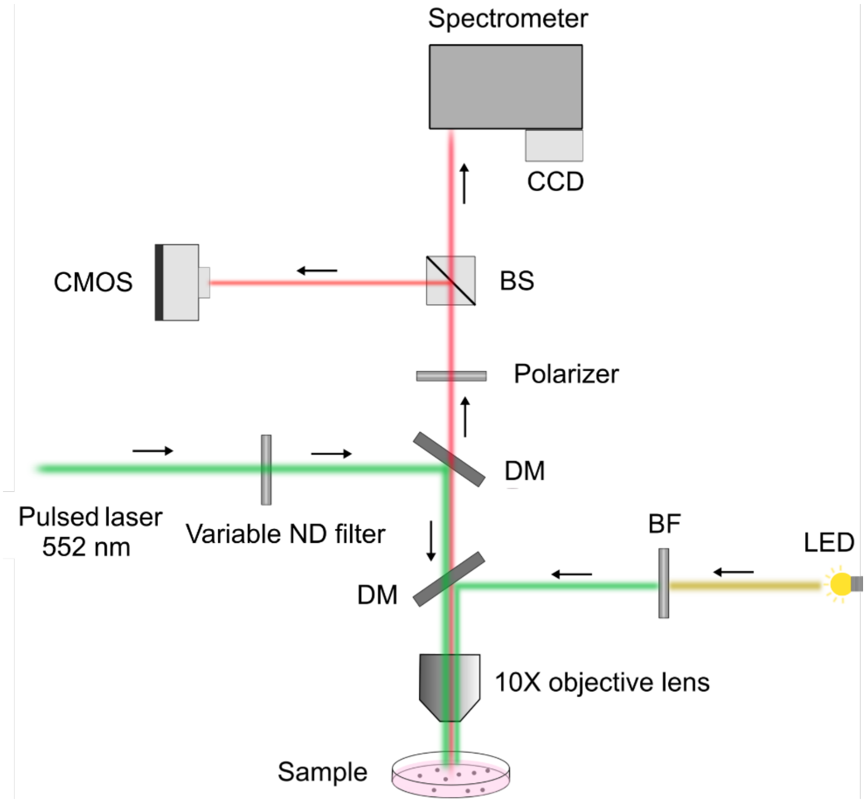


**Supplementary Figure 3**. Illustration of the experimental setup. The laser emission was split by a beam splitter and then propagated into the spectrometer and the CMOS camera, allowing the recording of the emission spectrum and transverse mode pattern. The LED excitation was used to observe the fluorescent images of the microlaser resonators. BS: beam splitter, DM: dichromic mirror, BF: bandpass filter, ND: neutral density.

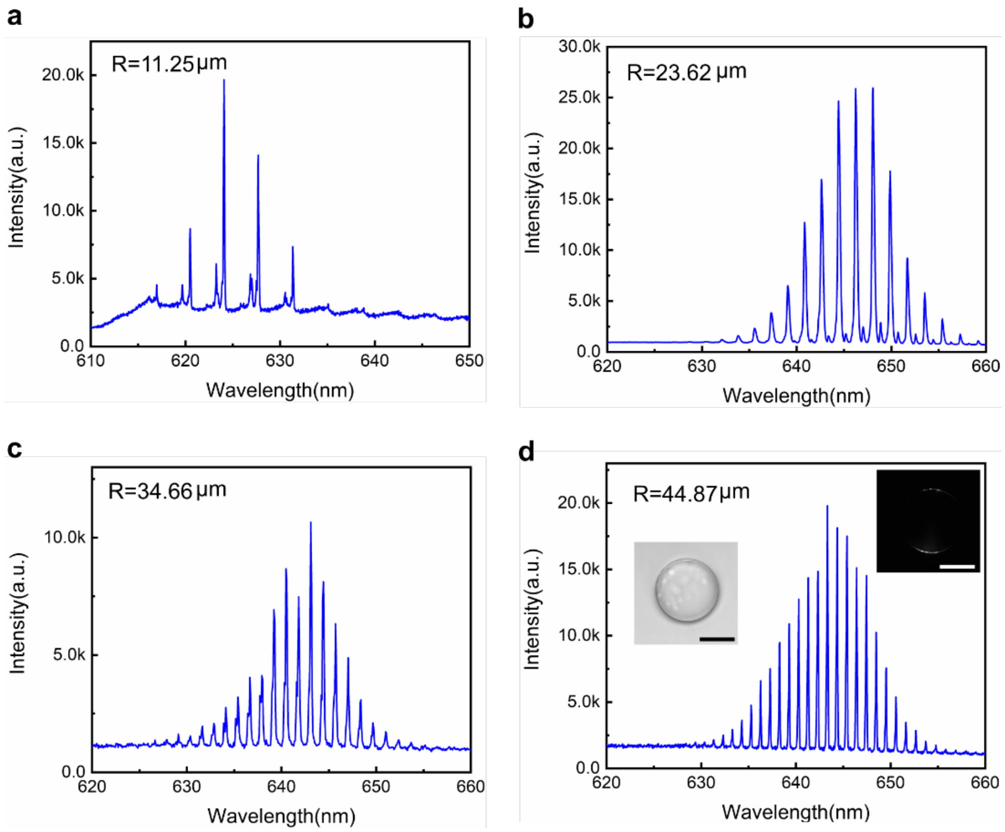


**Supplementary Figure 4**. (a)-(d) Emission spectra from different-sized microlasers. The smallest microlaser resonators that can generate laser can be around 11 μm (radius), showing fewer peaks and large FSR. With increasing the size, the laser peaks became more and the FSR became smaller. Inserts in (d) are the bright-filed image and WGM laser pattern of a microlaser with ~ 45 μm radius. Scale bar: 50 μm.

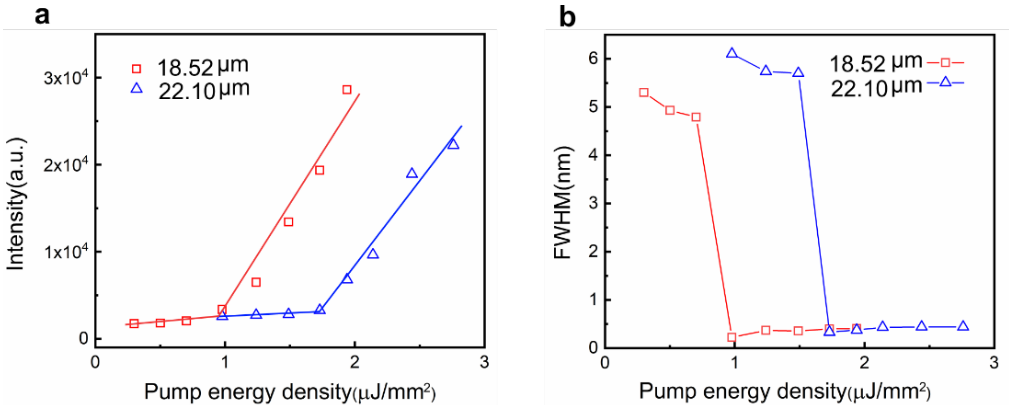


**Supplementary Figure 5**. (a)–(b) Laser threshold and FWHM characterization of the microlasers with a radius of 18.52 and 22.10 μm, respectively.

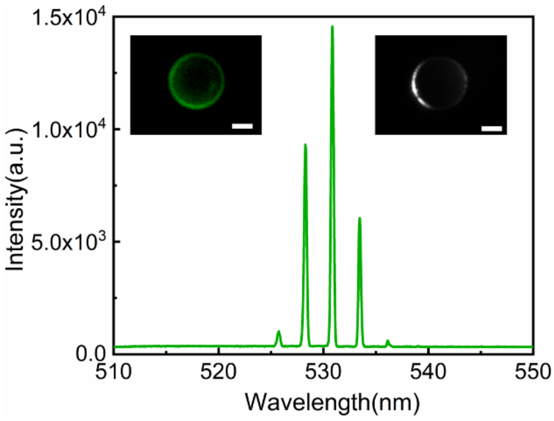


**Supplementary Figure 6**. Typical emission spectrum of a microlaser doped with Bodipy-1. Inserts are the fluorescence image (left) and the WGM laser pattern image (right). Scale bar: 10 μm.

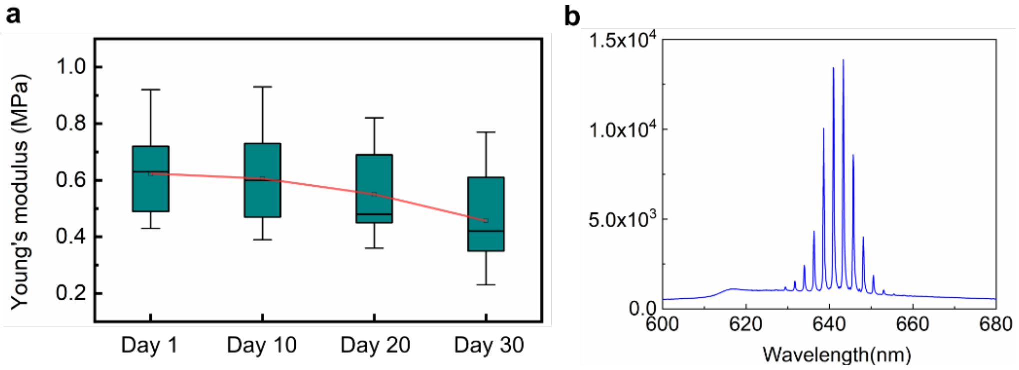


**Supplementary Figure 7**. (a) Young's modulus variation of the microlasers with time induced by degradation. (b) Emission spectrum of a microlaser after being stored in PBS for 30 days.

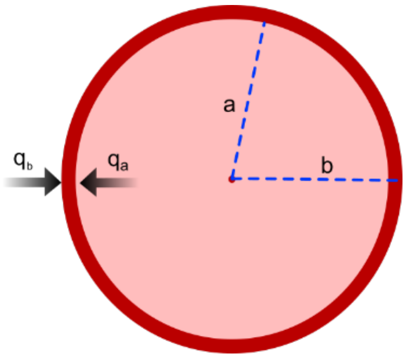


**Supplementary Figure 8**. Mechanical model of the hollow elastic microlaser. *a* and *b* are the inner radius and outer radius, respectively. $q_a$ and $q_b$ are inner stress and outer stress applied to the shell.

To theoretically calculate the radius displacement, we assume that the hollow microsphere is under a uniform force field. The inner radius is *a*, outer radius is *b*, the pressure applied on the inner surface and external surface is $q_a$ and $q_b$. According to the references[1, 2], when the microsphere is under stress, a point M (R, χ, φ) in the spherical coordinate system has the following equilibrium differential equation, geometric equation, and physical equation:

$$\frac{\partial\sigma_R}{\partial R}+\frac{2(\partial_R-\partial_T)}{R}+F_R=0(\rho\frac{\partial^2 u_R}{\partial t^2}) \quad (1)$$

$$\varepsilon_R=\frac{du_R}{dR},\varepsilon_R=\frac{u_R}{R} \quad (2)$$

$$\sigma_R=\frac{E}{1+v}\left(\frac{v}{1-2v}\theta+\varepsilon_R\right),\ \sigma_T=\frac{E}{1+v}\left(\frac{v}{1-2v}\theta+\varepsilon_T\right) \quad (3)$$

where $\theta=\varepsilon_R+2\varepsilon_T$, tangential strain $\varepsilon_R=\varepsilon_T$, tangential stress $\sigma_R=\sigma_T$, *E* is the elastic modulus or Young's modulus, *v* is Passion ratio, $F_R$ is the volumetric strain.
Then we could have

$$\sigma_R=\frac{E}{(1+v)(1-2v)}[(1-v)\varepsilon_R+2v\varepsilon_T],\ \sigma_T=\frac{E}{(1+v)(1-2v)}(\varepsilon_T+v\varepsilon_R) \quad (4)$$

Then we could have the governing differential equation for the displacement issues:

$$\frac{E(1-v)}{(1+v)(1-2v)}\left(\frac{\partial^2 u_R}{\partial R^2}+\frac{2}{R}\frac{\partial u_R}{\partial R}-\frac{2u_R}{R^2}\right)+F_R=0(\rho\frac{\partial^2 u_R}{\partial t^2}) \quad (5)$$

If we ignore the volumetric strain, the equation (5) can be simplified to

$$\frac{\partial^2 u_R}{\partial R^2}+\frac{2}{R}\frac{\partial u_R}{\partial R}-\frac{2u_R}{R^2}=0 \quad (6)$$

The equation (6) can be integrated to

$$u_R=AR+\frac{B}{R^2} \quad (7)$$

where *A* and *B* are integral constant that are determined by boundary conditions.
Then, combining equation (7) with equation (2) and (4), we could have the following equations:

$$\sigma_R=\frac{E}{1-2v}A-\frac{2E}{1+v}\frac{B}{R^3},\ \sigma_\theta=\frac{E}{1-2v}A-\frac{E}{1+v}\frac{B}{R^3} \quad (8)$$

The boundary conditions of microsphere are

$$(\sigma_R)_{R=a} = -q_a, \ (\sigma_R)_{R=b} = -q_b \tag{9}$$

Combining equation (8) and (9), we could have the A and B:

$$A = \frac{1-2v}{E} \cdot \frac{a^3 q_a - b^3 q_b}{b^3 - a^3}, \ B = \frac{1+v}{2E} \cdot \frac{a^3 b^3 (q_a - q_b)}{b^3 - a^3} \tag{10}$$

Thus, the radius displacement equation (7) can be given by

$$u_R = \frac{(1+v)R}{E}\left[\frac{\frac{b^3}{2R^3}+\frac{1-2v}{1+v}}{\frac{b^3}{a^3}-1} q_a - \frac{\frac{a^3}{2R^3}+\frac{1-2v}{1+v}}{1-\frac{a^3}{b^3}} q_b\right] \tag{11}$$

According to the FSR formula, the change of FSR is given by

$$\Delta FSR = \frac{\lambda^2}{2\pi} \frac{u}{b(b-u)} \tag{12}$$

Combine equations (11) and (12), we could get the relationship of the FSR change and applied stress.

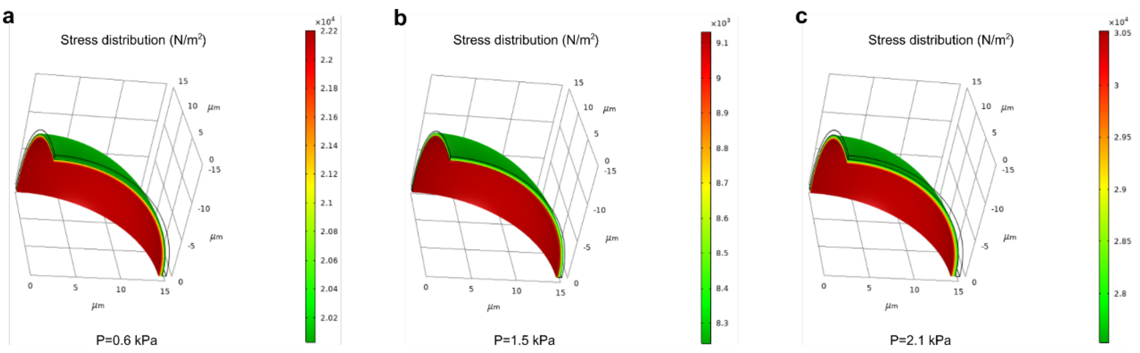


**Supplementary Figure 9**. (a)-(c) Simulated stress distribution on the microlasers under 0.6, 1.5 and 2.1 kPa, respectively. (1N $m^{-2}$ = 0.001 kPa)

**Table 1** Stress sensitivity versus the size of microlasers

| Radius (μm) | 12.5 | 15 | 17.5 | 20 | 25 | 30 |
|---|---|---|---|---|---|---|
| ΔFSR/ΔR (pm/μm) | 299.63 | 205.10 | 149.17 | 113.35 | 71.79 | 49.5 |
| ΔR/ΔP (μm/kPa) | 0.175 | 0.250 | 0.339 | 0.441 | 0.691 | 0.995 |
| Sensitivity (pm/kPa) | 52.44 | 51.28 | 50.57 | 50.02 | 49.58 | 49.25 |

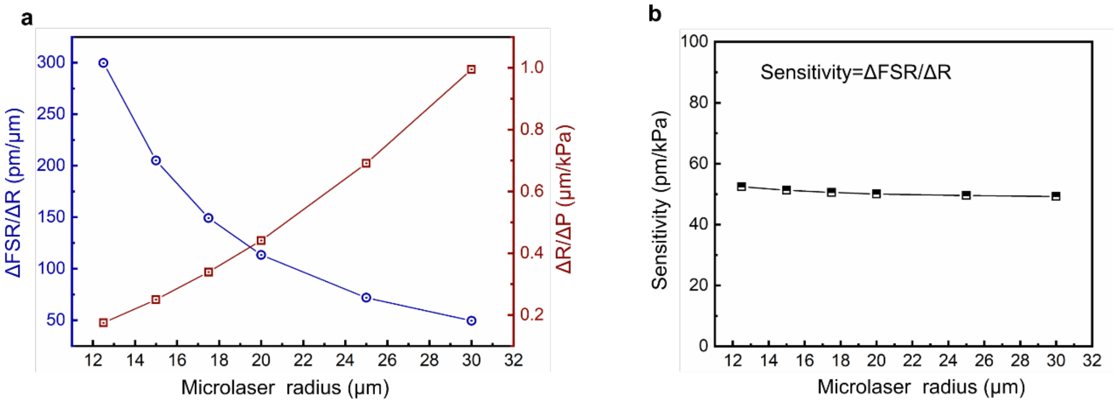


**Supplementary Figure 10**. (a) Ratio of FSR change to radius change and ratio of radius change to applied stress change under different microlaser size. (b) Sensitivity of microlaser under different sizes. This indicates that with the microlaser size increase, the microsphere deformation gradually increases under a certain stress. However, due to the small FSR of large microspheres, the FSR change decreases when a certain deformation happens. Thus, if we define the sensitivity of the microlaser as the ratio of FSR change to radius change, the sensitivity remain on a similar scale.

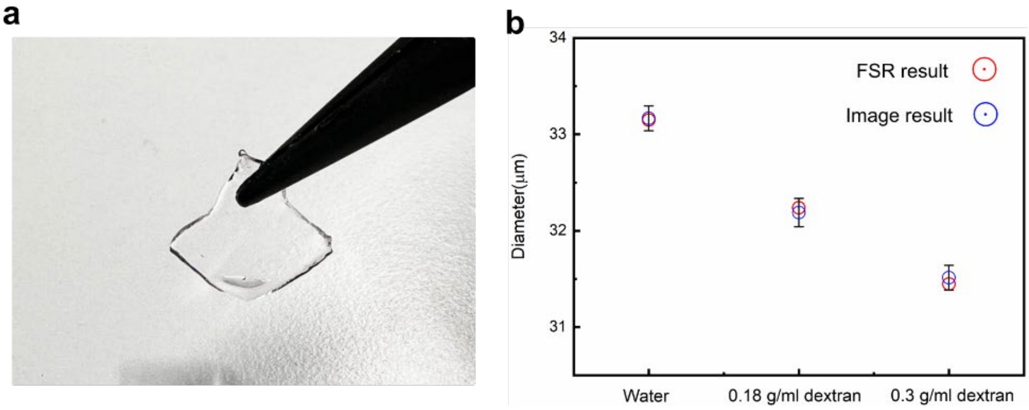


**Supplementary Figure 11**. (a) PAA hydrogel with microlasers embedded inside. The hydrogel is transparent, which enables both the measurement of fluorescent images and the laser spectrum collection. (2) Microlaser diameter measured by laser FSR and the fluorescent image, respectively. It is obvious that the diameter calculated by laser FSR agrees well with that derived from the fluorescent image.

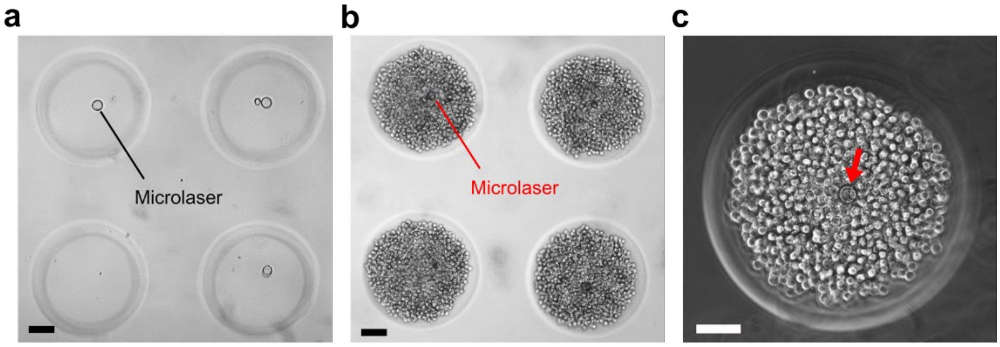


**Supplementary Figure 12**. (a) Microlasers seeded into the agarose microwell array. The concentration of the microlasers can be adjusted to ensure each microwell contained 1~2 microlasers. (b) Cells seeded into the microwells. (c) Enlarged view of a microwell that contained one microlaser and cells. Scale bar: 100 μm.

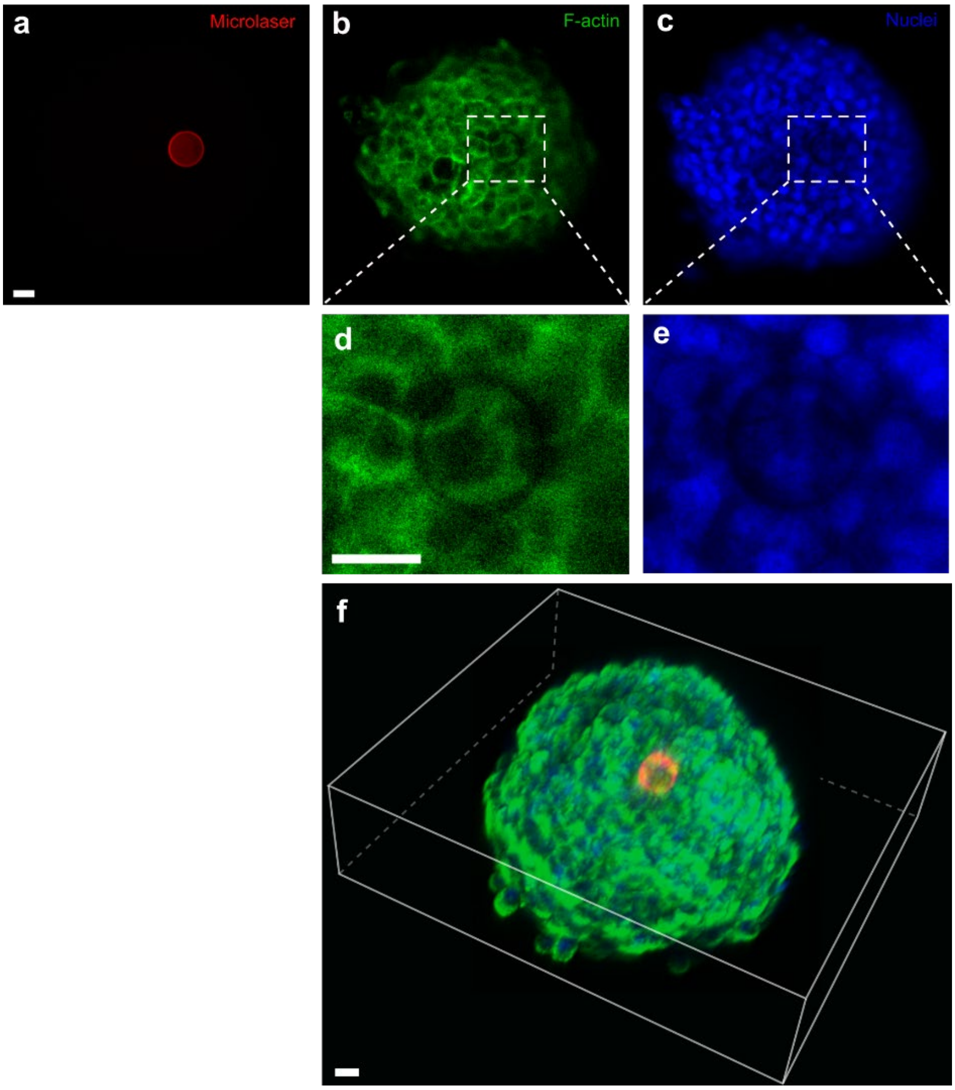


**Supplementary Figure 13**. Separated fluorescent channels of the image captured by the confocal microscopy. (a-c) Images of the microlases, F-actin and nuclei. (d-e) Enlarged view of the microlaser area. (f) 3D viewing of a multicellular spheroid with a microlaser embedded inside. Scale bar: 20 μm. The cellular structures can be observed through the microlasers. We hypothesized that this transparent hollow microspheres in the tissues could have an effect of optical lens, which can manipulate the light path[3, 4]. Thus, the illuminated cellular structures that near the microspheres can be projected inside the microspheres.

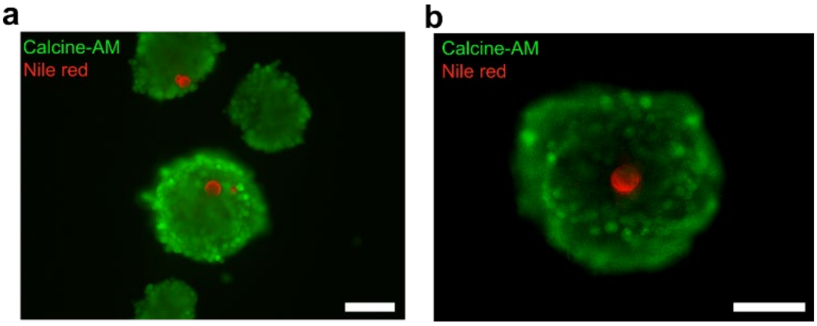


**Supplementary Figure 14**. (a)-(b) Cell viability characterization using Calcine-AM after co-culturing for 3 days. Scale bar: 100 μm. It indicated that the cells in the spheroids can maintain high viability when co-cultured with the microlasers for a long time.

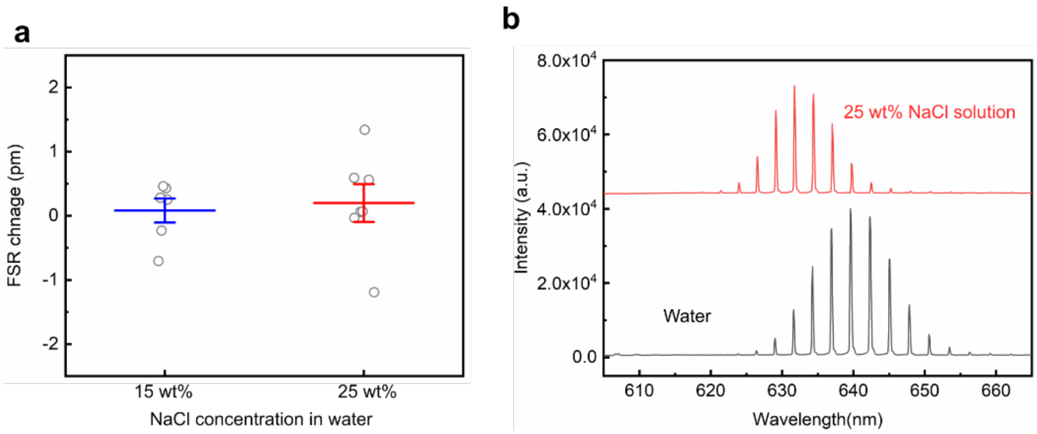


**Supplementary Figure 15**. (a) FSR change after the microlasers were immersed into the 15 wt% and 25 wt% NaCl solution. The refractive index of the 15 wt% and 25 wt% NaCl solutions are 1.364 and 1.378, respectively. This control experiment indicated that the refractive index of the surrounding environment has an ignorable effect on the microlaser FSR or the calculation of radius. (b) Laser spectra of a microlaser in water and in 25 wt% NaCl solution. Due to the change of the surrounding refractive index, the laser peaks showed several-nanometre blueshift. However, it didn't change the FSR.

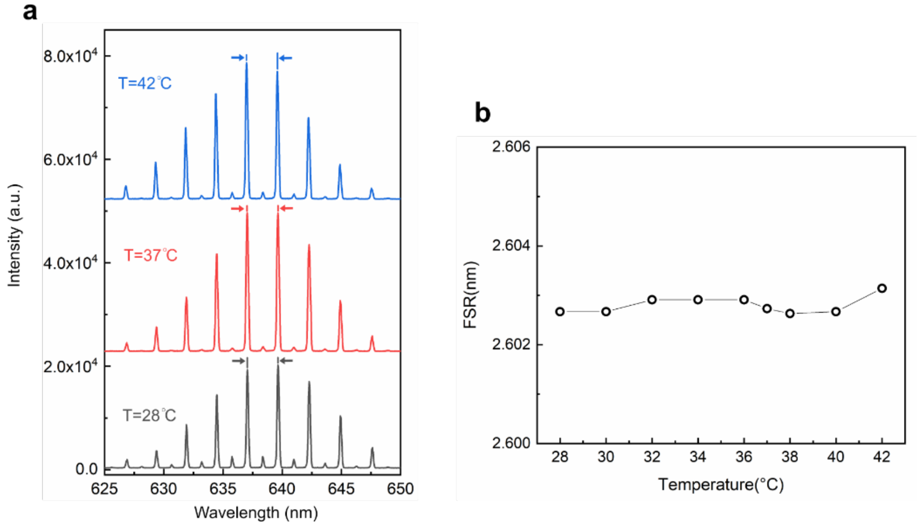


**Supplementary Figure 16**. (a) Emission spectra of individual microlaser at 28ºC, 37ºC and 42ºC, respectively. (b) FSR change under the temperature range from 28 ºC to 42 ºC. This control experiment indicates that in the biological experiments, the temperature fluctuation has no effect on the FSR, which will not influence the stress quantification.

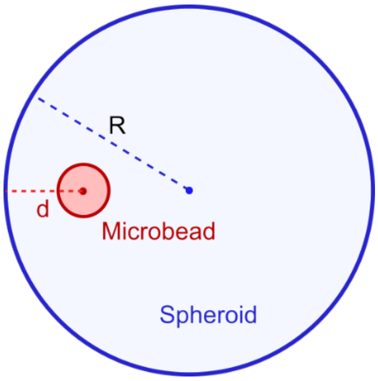


**Supplementary Figure 17**. Definition of the normalized distance (edge to centre). R: radius of spheroids, d: distance from the microlaser centre to the spheroid edge. The normalized distance = d/R.

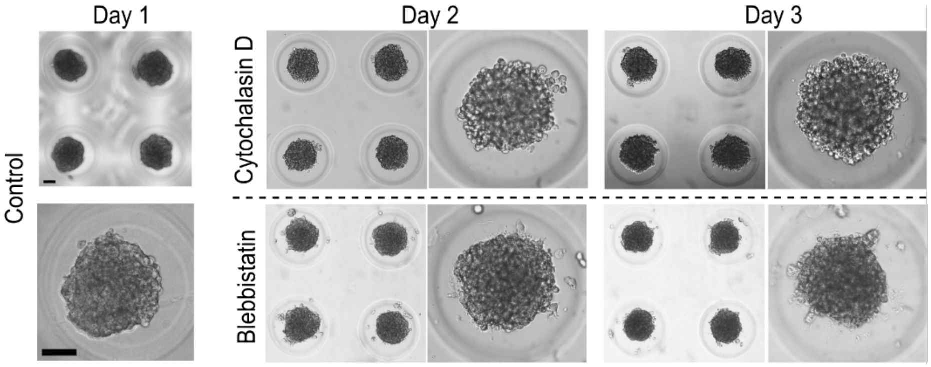


**Supplementary Figure 18**. A549 spheroid morphology before and after the treatment by Cytochalasin D and Blebbistatin. Scale bar: 100 µm. Before the treatment, the cells in spheroids were attached tightly and formed compact microspheres (control group). After the treatment by Cytochalasin D, the cells at the edge became round and unattached from the spheroids. In day 3, some cell debris could even be noticed at the edge. This resulted in the stress release inside the spheroids. In Blebbistatin, after the treatment, the cells started to be isolated from the spheroids. However, this was not very obvious.

**Supplementary video 1** Video-rate microscopy of the microlaser in multicellular tumor spheroids. (A) Lasing of a microlaser close to the surface of an A549 tumor spheroid under the bright-filed microscopy. The brilliant lasing indicated a high contrast to the bright-filed background. Scale bar: 100 µm. (B) Microlaser lasing in the tumor spheroid without the bright-field illumination. Scale bar: 100 µm. (C) Lasing of a microlaser located close to the centre of the spheroid. Scale bar: 50 µm.

**Supplementary video 2** Video-rate microscopy of the microlasers in small hESC-derived cardiac organoids. (A) A microlaser close to the surface of the organoid under bright-field microscopy. The compression and recovery can be observed. (B) A microlaser close to the centre of the cardiac organoid. Scale bar: 100 µm.

**Supplementary video 3** Video-rate microscopy of the microlasers in the large hESC-derived cardiac organoid. (A) A large hESC-derived cardiac organoid beating under bright-field microscopy. Scale bar: 200 µm. (B) Microlasers embedded inside the cardiac organoids under fluorescence microscopy. Scale bar: 200 µm. The fluorescent images of the microlasers are significantly blurry when deeply embedded. (C) Microlaser lasing in the large cardiac organoids. Scale bar: 200 µm.
